# Improved RNA stability estimation through Bayesian modeling reveals most *Salmonella* transcripts have sub-minute half-lives

**DOI:** 10.1101/2023.06.15.545072

**Authors:** Laura Jenniches, Charlotte Michaux, Linda Popella, Sarah Reichardt, Jörg Vogel, Alexander J. Westermann, Lars Barquist

## Abstract

RNA decay is a crucial mechanism for regulating gene expression in response to environmental stresses. In bacteria, RNA-binding proteins (RBPs) are known to be involved in post-transcriptional regulation, but their global impact on RNA half-lives has not been extensively studied. To shed light on the role of the major RBPs ProQ and CspC/E in maintaining RNA stability, we performed RNA sequencing of *Salmonella enterica* over a time course following treatment with the transcription initiation inhibitor rifampicin (RIF-seq) in the presence and absence of these RBPs. We developed a hierarchical Bayesian model that corrects for confounding factors in rifampicin RNA stability assays and enables us to identify differentially decaying transcripts transcriptome-wide. Our analysis revealed that the median RNA half-life in *Salmonella* in early stationary phase is less than 1 minute, a third of previous estimates. We found that over half of the 500 most long-lived transcripts are bound by at least one major RBP, suggesting a general role for RBPs in shaping the transcriptome. Integrating differential stability estimates with CLIP-seq revealed that approximately 30% of transcripts with ProQ binding sites and more than 40% with CspC/E binding sites in coding or 3’ untranslated regions decay differentially in the absence of the respective RBP. Analysis of differentially destabilized transcripts identified a role for ProQ in the oxidative stress response. Our findings provide new insights into post-transcriptional regulation by ProQ and CspC/E, and the importance of RBPs in regulating gene expression.

**Significance Statement:** Together with transcription and translation, RNA decay is one of the major processes governing protein production. Here, we have developed a new statistical approach that corrects for confounding effects when estimating RNA decay rates from RNA-seq in bacteria. Our more accurate decay rate estimates indicate that *Salmonella* transcripts have half-lives about three times shorter than previously thought. This approach allowed us to measure the effects of RNA-binding proteins (RBPs) on decay rates, identifying large cohorts of transcripts with changes in stability following RBP deletion and conditions where post-transcriptional regulation affects survival. Our method should lead to a reevaluation of RNA stability estimates across diverse bacteria and new insights into the role of RBPs in shaping the transcriptome.

## Introduction

Rapid adaptation of the proteome to environmental conditions is essential for the survival of microorganisms. RNA degradation is an important post-transcriptional process directly influencing protein abundance. The lifetime of bacterial RNA ranges from seconds to an hour (1) and depends on numerous factors, including transcript identity, genotype and growth condition (2). RNA-binding proteins (RBPs) in bacteria include structural components of the ribosome and global post-transcriptional regulators such as Hfq (3, 4) and CsrA (5) which play key roles in modulating translation and RNA stability in concert with a network of small RNAs (sRNAs) (6, 7). Beyond these model RBPs, recent years have seen the discovery of a menagerie of bacterial RBPs that bind hundreds or even thousands of transcripts (8–10), though their functions in shaping the transcriptome remain unclear.

In *Salmonella enterica* serovar Typhimurium (henceforth *Salmonella)*, these recently identified global RBPs include the less studied FinO-domain containing protein ProQ and the cold-shock proteins CspC and CspE. While individual deletion of any of these RBPs does not lead to clear growth phenotypes under standard laboratory conditions (11, 12), mounting evidence suggests they play important regulatory roles. ProQ has been shown to bind hundreds of mRNAs and sRNAs (13–15), affecting important biological processes including expression of virulence factors (12) and formation of antibiotic persisters (16). ProQ has also been shown to be involved in determining the outcome of regulatory interactions between some sRNAs and target transcripts (12, 15, 17), though many of the mechanistic details of these interactions remain unclear (18). CspC and CspE have been shown to play partially redundant roles in virulence, affecting survival in mice, motility, biofilm formation, and survival of bile stress (11, 19). The molecular details of how these RBPs affect phenotype are not clear, although at least some of the effects of ProQ and CspC/E are mediated by the direct modulation of mRNA stability. For instance, CspC/E have been shown to stabilize the mRNA of the bacteriolytic lipoprotein EcnB by blocking digestion by the endonuclease RNase E (11). ProQ on the other hand appears to preferentially bind 3^′^ UTRs where in a few cases it has been shown to protect transcripts from exonuclease activity (14, 20).

While these results provide hints at the mechanisms by which RBPs regulate target gene expression, in the absence of transcriptome-wide differential RNA stability measurements it remains unclear how common regulation through stability modulation is. A classical approach to study RNA stability is to halt transcription with the transcription initiation inhibitor rifampicin (21) and monitor RNA decay over time. This approach has been scaled to the whole transcriptome by combining it with microarrays (22, 23) and high-throughput sequencing (24). However, the presence of non-linear effects in the resulting time-course data makes inference of differences in decay rates between experimental conditions difficult.

RNA-seq analysis tools such as limma (25), edgeR (26), and DEseq (27) solve the problem of accurately estimating dispersion in experiments with many measurements but few replicates through an empirical Bayes approach (28). In empirical Bayes, information is pooled across transcripts under the assumption that transcripts with similar concentrations will exhibit similar biological and technical variation across samples, leading to more robust dispersion estimates. However, these tools are currently restricted to linear models, which limits the structure of models that can be implemented and hence the complexity of effects that can be considered. Recent progress in the optimization of sampling methods has made the development of fully Bayesian hierarchical models increasingly efficient and accessible. In particular, the Stan probabilistic programming language (29) separates model description from sampler implementation, allowing easy development and testing of complex hierarchical models. This provides a powerful framework for developing analysis methods for sequencing data that can accommodate complex experimental techniques.

Here, we investigate the effects of the undercharacterized RBPs ProQ and CspC/E on RNA stability across the entire transcriptome, starting from a fully Bayesian analysis of rifampicin treatment followed by RNA sequencing (RIF-seq). During model development, we discovered that accounting for confounding factors in stability assays conducted after rifampicin treatment dramatically affects the inferred half-life, leading us to substantially revise estimates for average mRNA half-life in *Salmonella* to less than 1 minute, compared to previous estimates in the range of 2 to 7 minutes in the closely related species *E. coli* (22–24). We develop a hypothesis testing procedure for determining differential decay rates that allows us to identify hundreds of gene transcripts destabilized in the absence of ProQ and CspC/E. We combine our differential stability estimates with other high-throughput datasets available for *Salmonella* to further characterize RBP interactions, identifying a role for ProQ in survival of oxidative stress. We additionally find a substantial population of long-lived transcripts that depend on RBPs for their stability, illustrating the importance of RBPs in shaping the bacterial transcriptome. Beyond its utility in investigating RBP interactions, our improved approach to determining transcript half-life suggests that RNA stability in bacteria has generally been overestimated and will need to be reassessed in other bacterial species.

## Results

### A progressive Bayesian analysis revises RNA half-lives

To determine transcriptome-wide half-lives under an infection-relevant condition, we applied RIF-seq to *Salmonella* at early stationary phase (ESP) where host invasion genes are expressed (30). Our RIF-seq workflow for data production and analysis is illustrated in **Figure 1A**: wild-type and isogenic RBP deletion strains were treated with rifampicin, and cellular RNA samples were collected over time to capture RNA decay dynamics. We collected data from eight time points following rifampicin treatment in three (Δ*cspC/E*), six (Δ*proQ*), or nine (wild-type) replicates (see Methods). We included ERCC RNA spike-ins (31) for normalization between samples. Additionally, we developed a center-mean normalization technique to remove batch effects between replicate samples (**Figure S2;** *Methods*). Subsequently, we fitted a Bayesian statistical model to the normalized data using Hamiltonian Monte Carlo with Stan (29).

**Figure 1.**
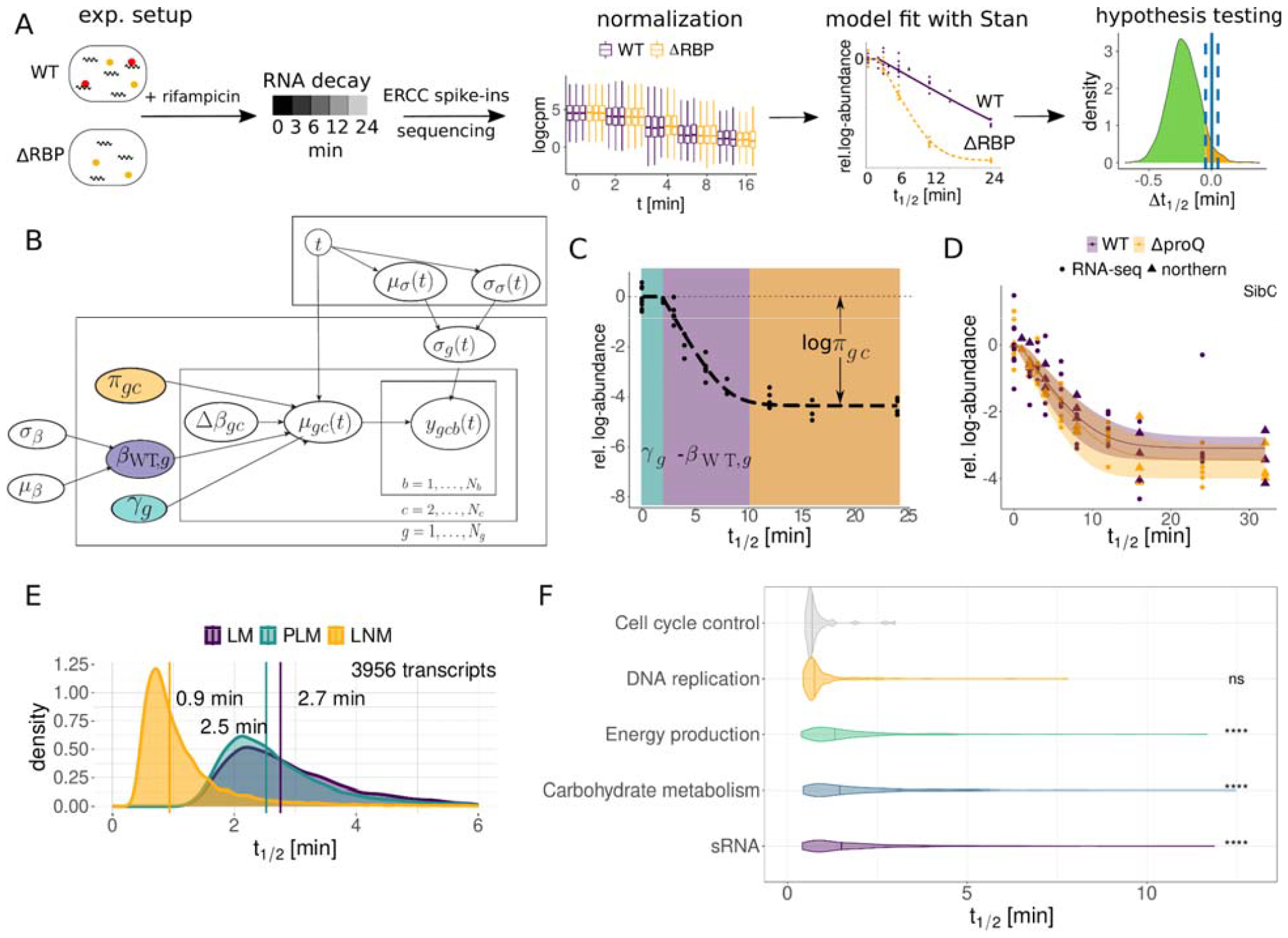
Pipeline and model description. (A) RIF-seq workflow: WT and ΔRBP strains are treated with rifampicin, cells are harvested at various time points and subjected to RNA-seq. Read counts are normalized before the extraction of biologically relevant parameters with a Bayesian model. Significant differences between strains are identified with Bayesian p values. (B) A plate diagram of the Bayesian models in this study. The layers indicate which indices and variables the parameters depend on. The LM is parametrized by the WT decay rate *β*_*WT,g*_ (purple). In the PLM, the gene-wise elongation time *γ*_*g*_ (green) is added. The LNM adds a baseline parameter *π*_*gc*_ (orange) which corresponds to the fraction of residual RNA (*π*_*gc*_ ∈ [0,0.2]). The WT decay rate *β*_*WT*_ is a gene-wise parameter that is modeled hierarchically and depends on the hyperparameters *μ*_*β*_ and *σ*_*β*_. The difference in decay rate *Δ β*_*gc*_ depends on the strain or condition *c*. The scale parameter *σ*_*g*_(*t*) captures variation by our decay model and depends on the time-dependent hyperparameters *μ*_*σ*_ (*t*) and *σ*_*σ*_ (*t*). (C) Representative example of a decay curve in the LNM, illustrating regimes dominated by the different model parameters. The period of transcription elongation *γ* is marked in green, the exponential decay with decay rate *β* in purple and the constant regime governed by the fraction of baseline RNA *π* in orange. (D) Comparison of RNA-seq and model fit with independent northern blot quantifications for SibC (13). (E) Hyperpriors and median of transcriptome-wide WT half-lives in the three Bayesian models. (F) Half-life distributions from the log-normal model for transcripts in selected COG categories. Adjusted p-values (compared to the first COG) were calculated using the Wilcoxon rank sum test.

We employed a progressive Bayesian workflow to arrive at our final model (**Figure 1B**). An advantage of Bayesian analysis is that it allows the modeler to formalize their beliefs about the data generating process and provides a variety of tools for model comparison and selection. In the case of RIF-seq data, the simplest expectation would be that RNA concentrations would exhibit a linear decay on a semilog scale, which could be fit by a simple linear model with gene- and condition-dependent decay rate *β*. While some of our observations met this expectation (**Figure S3A**), the vast majority of transcripts exhibited more complex dynamics that prevent accurate extraction of decay rates with a linear model (**Figure 1C; S3A**), leading to large unexplained variation at late time points (**Figure S3L**). To account for this, we introduced additional parameters that capture confounding effects in the data. The first confounding effect is a gene-dependent delay parameter *γ*, which captures the delay commonly observed in RIF-seq data before decay initiates (**Figure 1B&C**, in green). As has been previously described, this is due to ongoing transcription from RNA polymerase already bound to DNA, which rifampicin does not block (24, 32). The ongoing transcription compensates for decay, manifesting as a delayed decay, though our delay parameter may also capture unrelated effects such as the time needed for rifampicin to penetrate and act within cells. To support the relationship between ongoing transcription and the delay parameter, we performed an analysis of elongation times on 60 base sub-genic windows, finding a clear association between the estimated elongation time and distance to annotated transcription start sites (**Figure S3B**). We used this association to infer transcription rates from our data set (see **Figure S3C**, Methods) finding a median transcription rate of 22.2 nt/s (**Figure S3D**), comparable to previous estimates in *E. coli* (24).

The second confounding effect we corrected for was an apparent gene- and condition-dependent baseline RNA concentration *π* beyond which no further decay was observed (**Figure 1B&C**, orange). We were initially concerned that this effect may be an artifact of the pseudocount we used to avoid dividing by zero in our calculations; however, inspection of a number of decay curves illustrated that the observed baseline was generally well above the detection threshold (**Figure S3E**, see Methods). We also verified that the half-life of a transcript is generally constant along an operon (**Figure S3G**). In agreement with previous work (33) we found a small number of stable subregions which generally corresponded to known sRNAs (e.g. the FtsO sRNA excised from the *ftsI* mRNA, **Figure S3H**), but since this was not a general feature of transcripts we excluded this as a source of the observed baseline. To confirm that the baseline is not a result of our sequencing protocol, we used independent northern blot quantifications from a rifampicin treatment time course including late time points from previous studies (13, 14). These quantifications reproduced the observed baseline effect (**Figure 1D&S3I-J**), illustrating that this is a general feature of rifampicin RNA stability assays. Similar artifacts can be observed in published northern blot quantifications of rifampicin-treated *Salmonella* and *E. coli* in both ESP (34, 35) and during exponential growth (36). For wild-type *Salmonella*, we find that a median of 2.6% of the initial transcript concentration appears resistant to decay (**Figure S3K**), and that the exponential decay regime ends at different timepoints for different transcripts and genotypes (**Figure S6AB&C**). Whether this fraction is truly resistant to degradation or just degrades much slower than the rest of the transcript population is unclear. However, the median fraction of baseline RNA increases to 5.7% when *proQ* is overexpressed (**Figure S3K**), suggesting that nonspecific RBP-RNA interactions may play a role in degradation resistance.

To compare the models with and without these two confounding factors, we calculated the difference in the expected log pointwise predictive density (ELPD), a measure of the expected predictive accuracy of a model on out-of-sample data, using Pareto-smoothed importance sampling approximate leave-one-out cross-validation (PSIS-LOO, see Methods) (37). Comparing the difference in ELPD between a simple linear model, the piecewise linear model correcting only for extension time, and the full model (henceforth *log-normal* model) including the baseline correction showed a clear preference for the log-normal model, particularly at late timepoints (**Figure S3L&M**). Additionally, examination of the fitted variance unexplained by our decay model, σ_*g*_, illustrated the log-normal model captured the behavior of late timepoints better than the piecewise model (**Figure S3N-P**). Correcting for confounding factors has major implications for transcriptome-wide estimates of decay rates: while the linear and piecewise linear models produced median half-life estimates of 2.7 and 2.5 minutes, respectively, our final log-normal model estimates a median half-life of 0.9 minutes (**Figure 1E**).

To investigate whether transcripts encoding proteins involved in different cellular functions systematically differ in their stability, we calculated average half-lives across clusters of orthologous groups (COG) categories (38) (**Figure 1F; S5**). In agreement with previous work (22, 23), transcripts for genes involved in energy production and carbohydrate metabolism tended to be longer lived. We also found many sRNAs to have longer than average half-lives, with several of the longest-lived also having a connection to metabolism (e.g. GlmZ (39), Spot42 (40), and CyaR (41)). Among the least stable transcripts were those coding for genes involved in cell division (e.g. *ftsZ*) and DNA replication (e.g. *dnaA, dnaN*), suggesting tight control of their cognate proteins. Taken together, accurately modeling RNA-decay curves led to drastically reduced transcriptome-wide half-life estimates and allowed us to relate transcript stability to gene function.

### Steady-state abundance does not reflect changes in transcript half-life upon RBP deletion

To study the influence of the RBPs ProQ and CspC/E on transcript stability, we applied the log-normal model to our RIF-seq data for the *proQ* and *cspC/E* deletion strains, as well as a *proQ* overexpression strain (*proQ++*; see Methods). To prioritize transcripts with changes in stability, we developed a hypothesis testing procedure based on examination of the posterior distribution of the change in decay rate from the wild-type (**Figure S4A**) and estimated statistical significance by calculating Bayesian p-values (**Figure S4B**). Since Bayesian p-value distributions require calibration (42, 43), we used simulation studies to estimate the false discovery rate (FDR) (**Figure S4C-E**, see Methods). To evaluate our differential stability estimates, we examined known targets of ProQ (13, 14) and CspC/E (11) (**Figure 2A-C, S6A**). For deletion of ProQ we were able to confirm destabilization of the *cspD, cspE*, and *ompD* transcripts (**Figure 2A**), while the *cspC* transcript was hyperstabilized in the *proQ++* background (**Figure 2B**) in agreement with previous northern analysis (14). Of the mRNAs known to be stabilized by ProQ, we were only unable to detect destabilization for the *yfiD* transcript. Similarly, we found the *ecnB* transcript destabilized following *cspC/E* deletion (**Figure 2C**) as previously reported (11).

**Figure 2.**
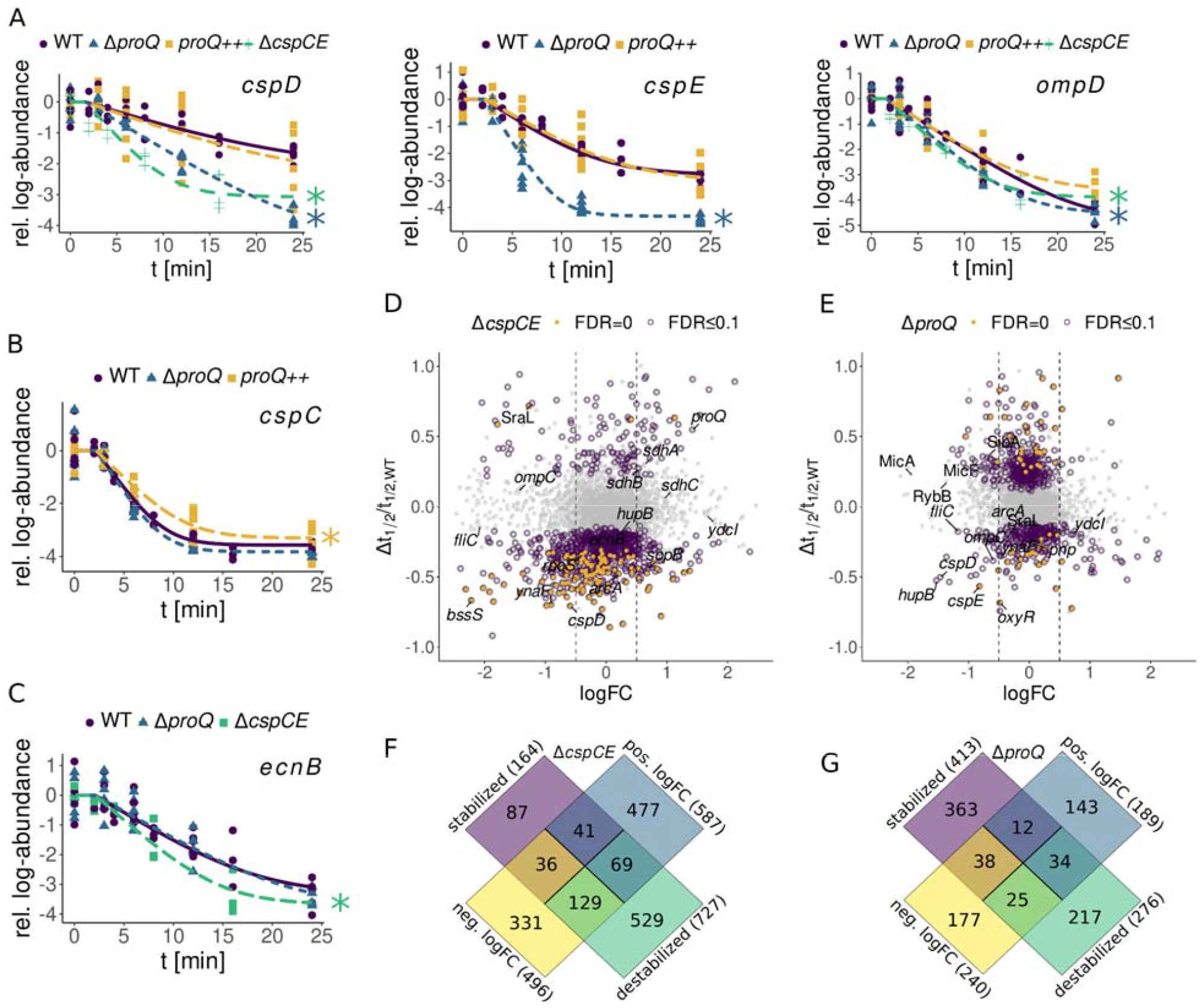
Differential analysis of transcript stability in the absence of CspC/CspE/ProQ. (A-C) Decay curves of all known ProQ (*cspD, cspE, ompD, cspC*) or CspC/E (*ecnB*) mRNA targets whose stability changes were confirmed by our study. See **Figure S6A** for sRNAs. Significant stability changes are marked with a star. (D) Relative difference in half-life vs. steady state log-fold changes between the Δ*cspCE* and the WT strain. (E) Relative difference in half-life vs. steady state log-fold changes between the Δ*proQ* and the WT strain. (F) Overlap between stability changes and steady-state log-fold changes in the Δ*cspCE* strains. (G) Overlap between stability changes and steady-state log-fold changes in the Δ*proQ* strain.

For both RBP deletions, we identify hundreds of transcripts with changes in stability at an FDR of 0.1 (**Figure 2D&E**). Deletion of *cspC/E*, whose role in maintaining transcript stability is less well explored, led to strong destabilization of a large cohort of transcripts (727), while only stabilizing 164 (**Figure 2F**). Curiously, we identified more transcripts which were significantly stabilized (413) than destabilized (276) following *proQ* deletion (**Figure 2G**), which was unexpected as prior studies have focused on ProQ’s stabilizing effect (13, 14). Nevertheless, stabilized transcripts tended to have much smaller changes in half-life, with a median change of 0.3 minutes (**Figure S6D**), compared to destabilized transcripts whose half-lives changed by 0.7 minutes on average. Overexpression of *proQ* restored stability for ∼95% of transcripts identified as destabilized by proQ deletion, but additionally significantly stabilized one fourth of the transcriptome (**Figure S6E&F**). The majority of the transcripts (> 90%) stabilized upon *proQ* overexpression were not affected by *proQ* deletion, suggesting these effects are largely non-specific.

A striking feature of our analysis of both strains was that changes in transcript half-life are not clearly related to changes in steady-state abundance upon RBP deletion (**Figure 2D-G**). In the *proQ* deletion strain, less than 10% of destabilized transcripts showed a statistically significant decrease in steady-state abundance. While this number was higher for the *cspC/E* deletion (∼18%), it was still only a minor fraction of the total number of destabilized transcripts. This might be explained by altered activity of other regulatory proteins. Deletion of either RBP led to perturbation of the stability of transcripts encoding major regulatory proteins including the anti-sigma factor Rsd, the transcription termination factors Rho and NusA, the alternative sigma factor RpoS, the nucleoid-associated HupA/B, and the cAMP receptor protein CRP (**Figure S6B**). For HupA/B and RpoS, we also observed reduced mRNA abundance in RBP deletion strains (**Figure S14**). Hence, loss of ProQ or CspC/E likely has complex, and in some cases indirect, effects on the global transcriptome. This suggests caution should be taken when deducing direct regulatory interactions from differential expression analysis of RBP-deletion mutants.

### Integrating high-throughput datasets identifies cohorts of mRNAs subject to known RBP regulatory mechanisms

The location of an RBP binding site within a transcript is often a key determinant of the mechanism of RBP regulation. To investigate the potential mechanisms underlying the stabilization activity of ProQ and CspC/E, we integrated our differential stability estimates with UV crosslinking and immunoprecipitation followed by RNA sequencing (CLIP-seq), which can localize RBP binding sites within a transcript. For ProQ, we reanalyzed an existing CLIP-seq dataset (14), identifying 833 peaks indicative of binding (see Methods). We produced new CLIP-seq datasets for both CspC and CspE and identified 1155 CspC and 861 CspE peaks, spread across 571 and 462 target transcripts, respectively (**Figure 3A&B;S7A-C**). In total, 717 transcripts are bound by at least one CSP, with 430 CspC peaks directly overlapping with a CspE peak (**Figure 3B**) supporting the previously reported partial redundancy between these proteins (11) and similar observations in *E. coli* (44). We saw especially dense clusters of CspC/E peaks in transcripts encoding genes involved in the TCA cycle, flagellar proteins, and proteins involved in host invasion associated with the *Salmonella* pathogenicity island 1 (SPI-1) type three secretion system (**Figure 3A**).

**Figure 3.**
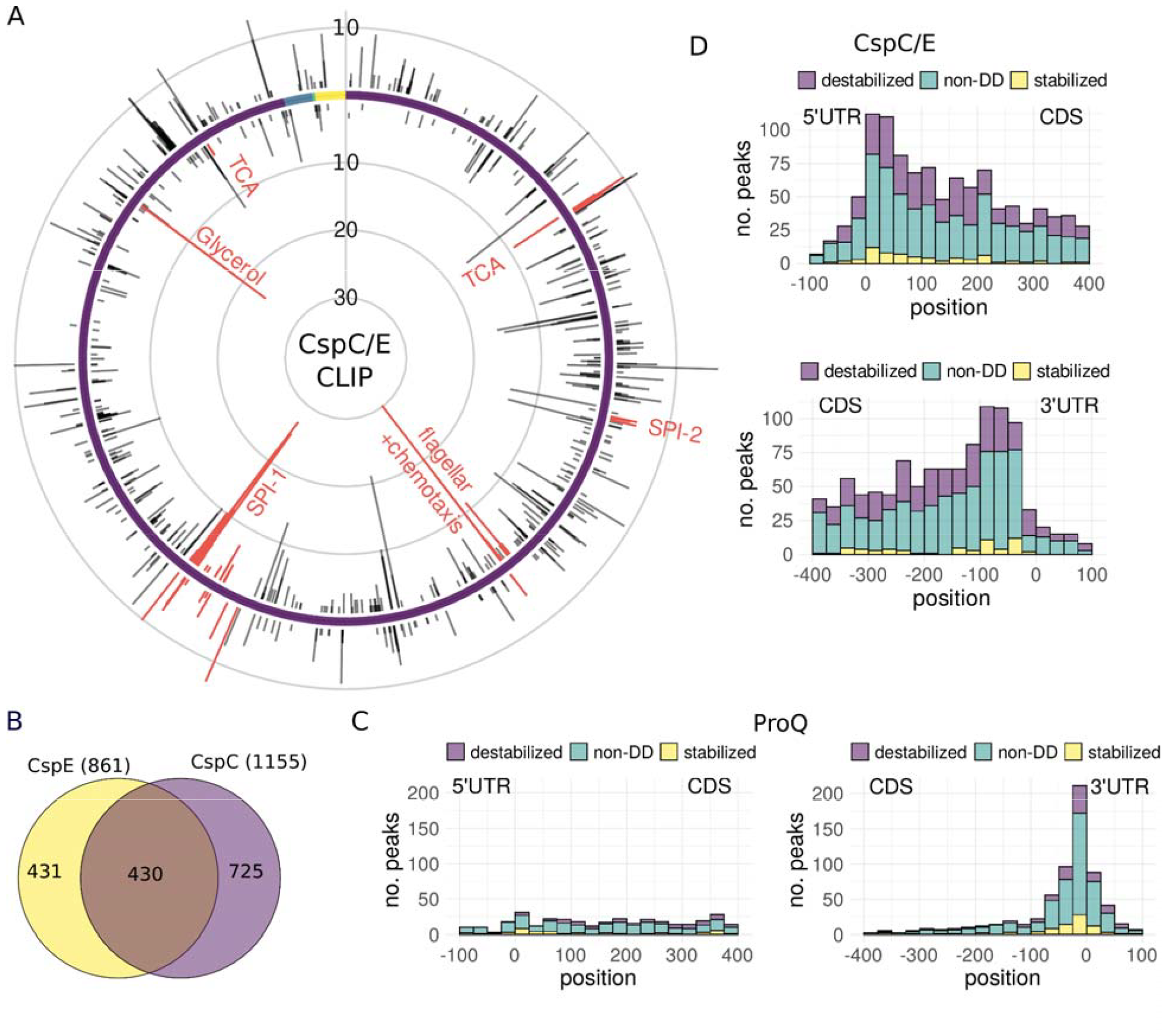
CspC/E CLIP-seq, comparison with RIF-seq results. CspC/E CLIP-seq analysis: (A) Number of CspC/E binding sites binned by genomic position for the positive (outer) and negative (inner) strand. The chromosome is indicated in purple and the three plasmids in blue, green and yellow. (B) Venn diagram of binding sites, with shared targets defined by an overlap by at least 12 bases between CspE and CspC sites. (C-D) Metagene plot of transcripts bound by the respective RBP, ordered by position of CLIP-seq peak relative to the start/stop codon. Target sequences are colored by the effect of RBP deletion on stability: destabilized (purple), stabilized (yellow), or no differential decay (non-DD, green).

We next examined the distribution of RBP binding sites across target transcripts, beginning with ProQ. As previously reported (14), ProQ binds predominantly at the end of coding sequences, with half of detected binding sites within 100 nucleotides of the stop codon (**Figure 3D;S7D**). Amongst those genes with 3′ binding sites, approximately 19% (86 transcripts) were significantly destabilized upon *proQ* deletion (**Table S3**). Besides the known interaction of ProQ with the *cspE* mRNA, these include transcripts encoding the SPI-1 effectors SopD and SopE2, involved in host cell invasion (45), and OxyR, a transcription factor involved in the oxidative stress response (46). The location of these binding sites suggests that ProQ may protect the 3′ ends of a large cohort of transcript from exoribonucleases attack, as previously shown for individual model transcripts (14, 20).

In contrast to ProQ, CspC and CspE binding sites were spread across coding (CDS) regions with only slight enrichment in the vicinity of the start and stop codons (**Figure 3C;S7D**). We identified 177 transcripts with a CspC and/or CspE binding site in the CDS or 5’UTR that were destabilized upon *cspC/E* deletion (**Table S4**). These included the *ecnB* transcript (**Figure S10A**), which has previously been shown to bind CspC and CspE *in vitro* and to be protected from RNase E by CspC/E *in vivo* (11). To further investigate the role of the CSPs in protection from RNase E cleavage, we combined our stability and CLIP-seq data with a published dataset mapping RNase E cleavage sites (47). We saw an enrichment of RNase E cleavage sites within CspC/E CLIP-seq peaks (410/2059 compared to a median of 331/2059 across 100 simulations, *p ≈* 0, see *Methods*), but the majority of CspC/E binding sites did not directly occlude known RNase E cleavage sites. Furthermore, the presence of an RNase E cleavage site within a peak did not appear to influence differential decay rates upon *cspC/E* deletion (**Figure S7E**). This suggests that rather than directly protecting cleavage sites, CspC/E may interfere with RNase E scanning (48). This is further supported by the fact that destabilized transcripts have a median of two CspC/E binding sites, while ligands without stability changes have a median of one binding site (**Figure S7F**), suggesting multiple CspC/E proteins must bind to create an obstruction of sufficient size to interfere with RNase E scanning (49).

### RBPs play overlapping and complementary roles in infection-relevant pathways

To investigate the physiological consequences of RBP deletion, we identified pathways enriched in differentially stabilized and differentially expressed transcripts in the *proQ* and *cspC/E* deletion strains with the GSEA algorithm (50) (**Figure 4A & S8A**). Intriguingly, we found a large overlap in enriched gene sets in both deletion backgrounds that may in part reflect destabilization of the *cspE* mRNA in the *proQ* deletion background (14). On the level of stability this included responses to extracellular stimulus and oxidative stress, flagellar assembly, and metabolite transport and utilization pathways including the phosphotransferase system and glyoxylate and dicarboxylate metabolism. Several of these gene sets were also enriched in differentially expressed transcripts, though the directions of the changes were often inconsistent with the observed effects on stability. For instance, genes involved in flagellar assembly were expressed at lower levels in both deletion strains despite their transcripts being stabilized (**Figure 4A, S8B & S12**). Some pathways, such as aerobic and anaerobic respiration, showed consistent changes in expression levels across both strains despite no clear shared enrichment on the level of stability.

**Figure 4.**
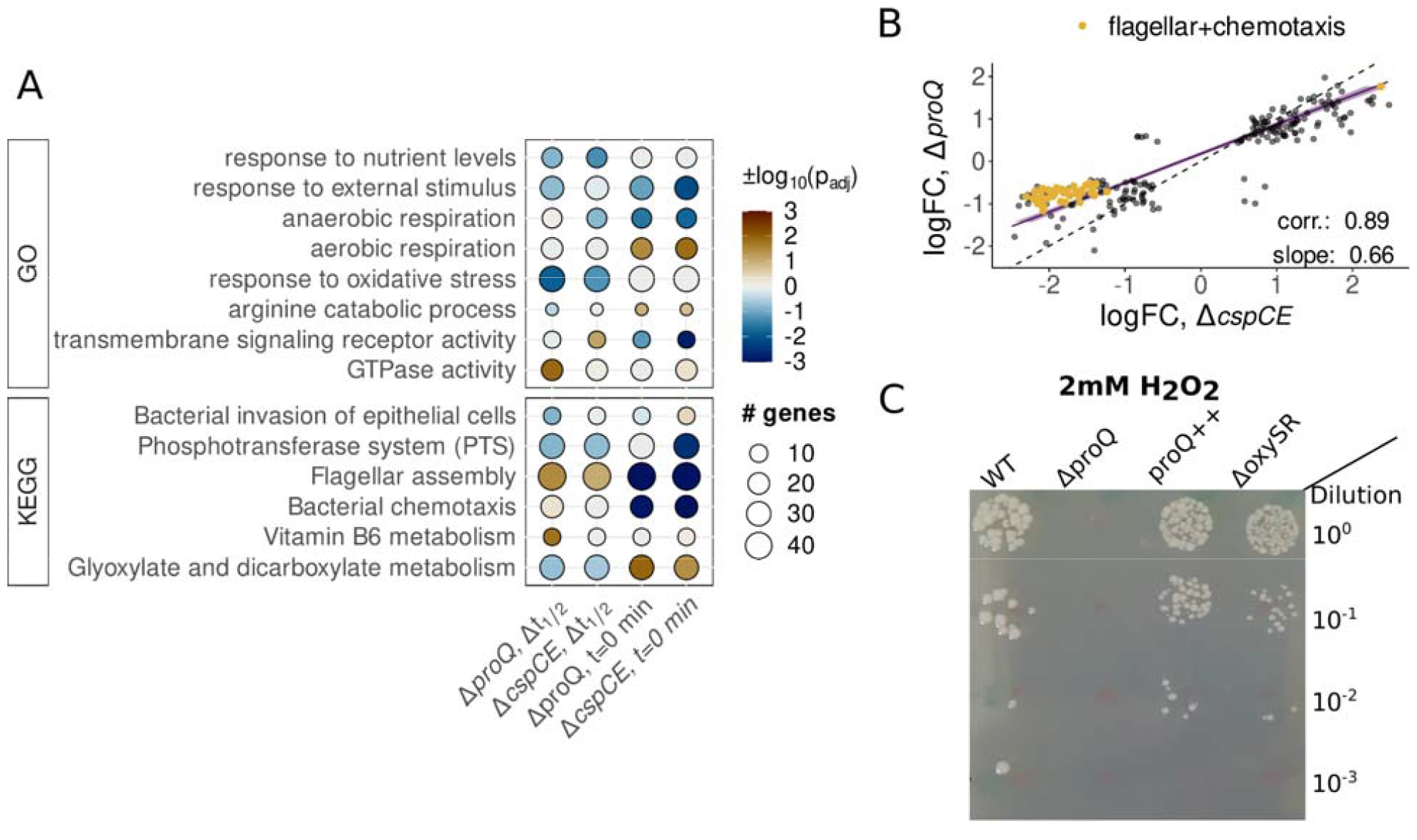
Integrative analysis of RBP binding and transcript stability. (A) Comparative analysis of pathways enriched in cspCE and proQ RIF-seq data. Pathways enriched in transcripts destabilized or with negative steady-state log-fold changes (min) upon RBP deletion are marked blue. Pathways enriched in stabilized transcripts or positive log-fold changes are marked brown. (B) Genetic features with significant log-fold changes in both the *proQ* and the *cspC/E* deletion mutant. (C) Exposure of Δ*proQ, proQ*++ and Δ*oxySR Salmonella* strains to 2mM of hydrogen peroxide.

The large overlap in pathways affected at the level of stability and expression between the two RBP deletion strains led us to investigate the relationship between ProQ- and CspC/E-mediated regulation. We examined transcripts significantly differentially expressed in both strains, finding a strong correlation between the steady-state log fold-changes (**Figure 4B**, r=0.89). The slope of a line fitted to these changes indicated stronger average changes in the *cspC/E* deletion; this was particularly clear for genes involved in flagellar assembly and chemotaxis which exhibited an ∼2-fold lower expression in the Δ*cspC/E* background compared to Δ*proQ*. The lack of a similarly strong correlation for changes in transcript stability (**Figure S8C & D**) suggested that some of the similarities in changes in steady-state mRNA abundance between the two deletion strains may be due to indirect regulation, with that in the Δ*proQ* background possibly mediated by changes in CspE expression.

Despite the large overlaps in mRNA stability and abundance changes, there were a number of changes specific to each RBP, though these were often in the same pathways. For instance, deletion of each RBP affected the stability of a discrete set of secreted effectors involved in host cell invasion (**Figure S9D**). We also identified changes for transcripts involved in *porin activity* (**Figure S13**), which may be related to ProQ’s originally described role in osmoregulation (51, 52). We found a ProQ CLIP-seq peak in the 5’ UTR of the *proP* transcript, but this did not appear to affect transcript stability or abundance, supporting a potential role in regulation of translation (53). Both *proQ* and *cspC/E* deletion affected stability and expression of the transcript for the major porin OmpD (**Figure 2A**), while effects on the transcripts of the well-characterized osmolarity-responsive OmpF and OmpC appeared to be RBP-specific (**Figure S13**). Another pathway with changes upon *proQ* and *cspC/E* deletion was in the oxidative stress response, where we saw a stronger enrichment for destabilized transcripts in the Δ*proQ* strain (**Figure 4A & S11**). Transcripts destabilized by *proQ* deletion included those encoding for the oxidative stress regulator OxyR (**Figure S9B & C**), the superoxide dismutase SodB, the catalase-peroxidase KatG, and the DNA protection during starvation protein Dps; however, few of these transcripts showed significant differences in mRNA abundance.

To test if destabilization was predictive of phenotype, we exposed the Δ*proQ* and *proQ*++ strains to varying concentrations of hydrogen peroxide, including a Δ*oxyR/S* strain as a control. After exposure to 1.5mM H_2_O_2_ we saw a survival defect for both Δ*proQ* and *proQ*++ strains intermediate between wild-type survival and that of the Δ*oxyR/S* strain (**Figure S9D**). However, this defect was concentration dependent: exposure to 2mM H_2_O_2_ led to a severe survival defect in the Δ*proQ* strain that could be complemented by *proQ* overexpression (**Figure 4C**), while Δ*oxyR/S* behaved similarly to wild-type. This indicates that the Δ*proQ* survival defect after exposure to high concentrations of H_2_O_2_ is independent of any effects ProQ has on the stability of the *oxyR* transcript and likely depends on the effects of ProQ on other transcripts involved in the oxidative stress response. The sensitivity of the Δ*proQ* strain to oxidative stress also shows that changes in transcript stability can be predictive of RBP deletion phenotype, even without corresponding changes in transcript abundance under standard growth conditions.

### Long-lived transcripts rely on global RBPs for their stability

To further examine the global impact of RBPs in shaping the transcriptome, we investigated the relationship between our wild-type half-life estimates and RBP binding as determined by CLIP-seq for five major *Salmonella* RBPs at early stationary phase: ProQ (14), CspC/E (this study), Hfq and CsrA (54). After sorting transcripts by stability, we saw a clear association between wild-type half-life and RBP binding, with over half of the 500 most stable transcripts (*t*_*1/2*_ > 2.5 min) bound by at least one RBP (**Figure 5A & S9E**). While the probability of detecting a CLIP-seq peak increases with transcript abundance (**Figure S9F&G**), we found no correlation between transcript abundance and stability (**Figure S3F**) suggesting the relationship between RBP binding and stability is unlikely to be an artifact of our measurements. Long-lived transcripts are also more likely to be destabilized upon RBP deletion than shorter-lived ones, regardless of RBP-binding. Of the 500 most stable transcripts, 32% are significantly destabilized in the absence of ProQ and 51% in the absence of CspC/E. Investigating the relationship between transcript half-life and differential half-life upon RBP deletion revealed large changes in median half-life for stable transcripts (**Figure 5B**), indicating that long-lived transcripts are not only bound by RBPs but also rely on them for their stability.

**Figure 5.**
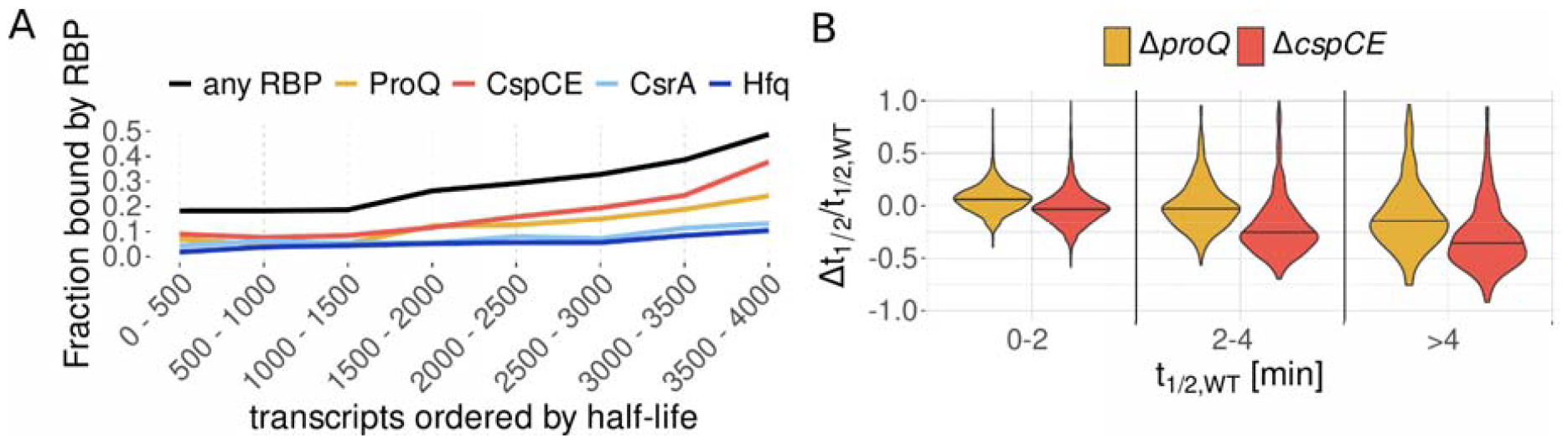
Global effect of RBP binding on transcript stability. (A) Fraction of transcripts bound by RBPs. Transcripts were ordered by half-life. The distribution of half-lives for each individual group can be found in **Fig. S9E**. (B) Relative change in half-life in the Δ*proQ* and Δ*cspCE* strain for transcripts with a WT half-life of 0-2 min, 2-4 min, and greater than 4 min.

## Discussion

Together with transcription and translation, mRNA degradation is one of the fundamental processes controlling protein production in the cell. Rapid turnover of mRNAs underlies the ability of bacteria to rapidly adapt to new conditions: as protein production is constrained by the translational capacity of the available ribosome pool (55), clearance of transcripts encoding for unneeded proteins is essential to change the composition of the proteome. Previous work based on RNA-seq and microarray analysis of rifampicin time course data in *E. coli* and *Salmonella* has reported average mRNA half-lives in the range of 2 to 7 minutes (22–24, 56– 58). The most similar prior RNA-seq study to our own reported an average half-life of 3.1 minutes across ∼1200 transcripts in *E. coli* grown to stationary phase (24), over three times our estimated average decay rate of 0.9 minutes in *Salmonella* at ESP. This discrepancy appears to originate from not accounting for the baseline stable RNA concentration, leading to a systematic underestimation of the decay rate. Interestingly, our estimates are in the range of those derived from classic experiments that pulse radiolabeled bulk RNA and determined average mRNA half-life to be ∼0.7 minutes in exponentially growing *E. coli* (59), far shorter than any other subsequent estimates based on high-throughput approaches. This rapid decay may in part underlie the ability of bacteria to rapidly adapt their transcriptomes, as constant transcription would be required to maintain mRNA concentrations. Across bacteria, mRNA half-lives have primarily been determined by rifampicin treatment followed by sequencing or microarray analysis in those bacteria where transcriptome-wide measurements are available (60) and appear to contain similar artifacts to those observed here. For instance, biphasic decay curves suggestive of a stable baseline transcript fraction have also been observed in the slow-growing *Mycobacterium smegmatis* (61). Our results thus suggest that mRNA stability has likely been widely overestimated and that a general reevaluation of bacterial transcript stability is in order.

Our hierarchical Bayesian analysis of RIF-seq data provides a principled framework for the analysis of RNA turnover, including the determination of differential decay rates after deletion of an RBP of interest. The flexibility of Bayesian analysis allowed us to account for nonlinearities due to confounding factors like transcription elongation after rifampicin addition and RNA baseline concentration, removing substantial biases in our determination of decay rates. Despite our best efforts, it is likely that there are still some limitations to our analysis. For instance, our control for false discovery rates means that we have likely missed some genuine instances of differential decay. Our simulations suggest an 85% sensitivity for the most precisely measured transcripts, but this falls to ∼30% when considering the whole transcriptome (**Figure S4D**). Other limitations may be due to uncontrollable effects in the data. For example, some *Salmonella* promoters have previously been shown to respond specifically to subinhibitory rifampicin (62), which could introduce some bias to decay rate estimates for affected transcripts should similar effects occur with the rifampicin concentrations used here. Manual inspection of our decay curves suggests this is unlikely to be a widespread problem in our data. Similarly, if RBP deletion leads to modulation of expression of cellular RNases, our individual differential decay rates may not be reflective of the differential decay induced by simple ablation of an RBP binding site. Such a bias dependent on the cellular context of rifampicin treatment has previously been observed for the sRNA RyhB whose stability critically depends on the presence of its target mRNAs (63).

Regardless of potential biases, high-throughput methods provide at least one major advantage over classical molecular approaches to RBP characterization: numbers. While our approach does not definitively demonstrate a causal link between RBP binding and transcript fate, we are able to provide high-confidence targets for future molecular characterization. Where previously a small handful of transcripts were known to be stabilized by 3′ binding of ProQ in *Salmonella*, we find 86 candidates. Similarly, we expand the number of transcripts known to be stabilized by CspC/E from two to a predicted cohort of 177. By combining CLIP-seq (14) and RNase E cleavage profiling (47) with our differential stability data, we have defined cohorts of transcripts likely subject to particular modes of RBP regulation. Depending on the binding site within a transcript, up to 44% (CspC/E, CDS) and 32% (ProQ, close to the stop codon) of direct RBP targets showed altered stability upon deletion of the respective RBP. However, in both cases transcripts stabilized by RBP binding are outnumbered by those apparently bound, but unaffected at the level of stability, raising numerous questions about RBP function. How are transcripts stabilized by RBPs differentiated from those that are not? Do RBP interactions that do not affect stability perform other functions in the cell? Our analysis suggests CspC/E may protect some transcripts from RNase E through a roadblock mechanism (48) which could be investigated further by mapping Rnase E cleavage sites in the presence and absence of CspC or CspE. (48) Additionally or alternatively, CspC/E targets may be regulated at the level of translation (64) or antitermination (65) through the manipulation of mRNA secondary structure as has been shown for the targets of other CSPs. Alternative roles of ProQ remain to be well defined, but it has been shown to play a role in gene regulation by sRNAs (12, 17). By defining and partially characterizing RBP targets, our data provides a starting point for the molecular investigations needed to further define the functions of CspC/E and ProQ.

The degree to which post-transcriptional regulation shapes the bacterial proteome has long been controversial. Recent work has suggested that, on average, protein concentrations are primarily determined by promoter on rates with post-transcriptional regulation playing only a minor role (66). Here in contrast, we have shown that deletion of bacterial RBPs thought to act primarily at the post-transcriptional level leads to large changes in both RNA stability and steady-state transcript concentration, and strong phenotypes have been observed for RBP deletion in a variety of conditions (5, 11, 67). How can these findings be reconciled? Our data provides at least two potential answers. First, as suggested by the effects of *proQ* and *cspC/E* deletion on the stability of various global regulators (Figure S6B&S14), modulation of stability or translation of single transcriptional regulators may ultimately cause phenotypic changes by indirectly affecting the promoter on rates of a large cohort of transcripts. The lack of correlation we observe between changes in steady-state RNA levels and differential stability (**Figure 2D&E**) indicates that such indirect effects are widespread. Secondly, our analysis shows that the majority of RNA half-lives are concentrated at less than 1 minute (**Figure 1E**), and it is indeed difficult to understand how further destabilization through post-transcriptional regulation could have strong effects on translation. However, the half-life distribution is long tailed, with ∼500 transcripts having half-lives of greater than 2.5 minutes and being preferentially bound by RBPs (**Figure 4A**). The stability of this population of transcripts is strongly affected by RBP deletion (**Figure 4B**), further suggesting they may be the major targets of post-transcriptional regulation.

An accumulating body of work suggests that the post-transcriptional regulatory networks scaffolded by RBPs are interconnected. At least two Hfq-dependent sRNAs also serve as sponges for CsrA (68, 69), and RNA-RNA interactome studies have observed a substantial fraction of shared targets between Hfq and ProQ (15). Regulatory interactions between cold shock proteins (CSP) have long been observed, with deletion of particular CSPs leading to the induction of others (11, 70), presumably through undescribed feedback mechanisms. The *cspE* mRNA has previously been used as a model for understanding the molecular mechanism of ProQ protection of 3′ ends (14); our results suggest some fraction of the change in steady-state transcript levels observed in the *proQ* deletion strain may be the result of indirect regulation through CspE (**Figure 5B**). Additionally, both RBPs affect the stability of mRNAs in similar pathways (**Figure 5A**), though often by targeting different transcripts, as for the SPI-1 effectors (**Figure S8B**). We also find effects for both strains on the stability of the CsrA-sponging sRNA CsrB, with *proQ* and *cspC/E* deletion having opposite effects on half-life (**Figure S10B & C**), adding a further potential connection between RBP regulatory networks. Our reanalysis of publicly available CLIP-seq data suggests that a substantial number of mRNAs are targeted by two or more RBPs (**Figure S10D**). What this apparently dense interconnection between RBP-mediated regulatory networks means for the cell, and how RBP activity is coordinated to maintain homeostasis in diverse environmental conditions, is an open question that will likely take significant conceptual advances to answer.

## Data Availability

Data deposition: All sequencing data reported in this paper have been deposited in the Gene Expression Omnibus (GEO) database, https://www.ncbi.nlm.nih.gov/geo (SuperSeries no. GSE234010). Transcript annotations and source code for the Stan models have been made available at https://github.com/BarquistLab/RIF-seq_repo

## Acknowledgements

We thank Joel Belasco, Erik Holmqvist, Susan Gottesman and Anke Sparmann for insightful comments on the manuscript, and Alexandre Smirnov for providing northern blot quantifications from (13). This project was funded in part by the Bavarian State Ministry for Science and the Arts through the research network bayresq.net (to LB, JV).

## Methods

### Media and growth conditions

For all experiments in this study, broth cultures were grown from single colonies overnight at 37 °C in LB medium (5 g/L of yeast extract, 5 g/L of NaCl, and 10 g/L of Tryptone/Peptone ex casein; Roth). Subsequently, cultures were diluted 1:100 in fresh medium, and further grown at 37°C with shaking at 220 rpm to an OD600 of 2.0 (early stationary phase (ESP), a SPI-1 inducing condition (30)).

### Bacterial strains and plasmids

*Salmonella enterica* serovar Typhimurium strain SL1344 (strain JVS-1574 (71)) is considered wild-type (WT). The generation of *proQ* and *cspC/E* deletion strains by lambda red homologous recombination (72) has been previously described (11, 13). For the *proQ*++ strain, a strain containing plasmid pZE12-ProQ was used as previously described (12, 13). The complete lists of bacterial strains, plasmids, oligos and antibodies used in this study are provided in SI Appendix, Table S1-4.

### Rifampicin assay protocol for sequencing

Wild-type (WT), ΔRBP and RBP++ strains were grown until an OD_600_ of 2.0 in three (WT, *cspCE*) or six (WT, *proQ, proQ*++) replicates. The cultures were treated with 500μl/ml of rifampicin (stock solution 50mg/ml resuspended in DMSO). Samples were taken before (*t* = 0min) and after 3, 6, 12, 24 min (proQ) or 2, 4, 8 and 16 min (cspCE) of rifampicin treatment. 2ml were collected for each sample, immediately mixed with 20% vol. stop mix (95% ethanol, 5% phenol) and snap frozen in liquid nitrogen.

Subsequently, the samples were thawed on ice and centrifuged for 20 min at 4500 rpm. Bacterial pellets were resuspended in 600 μl of 0.5 mg/ml of lysozyme in TE buffer pH 8 and transferred into a 2 ml Eppendorf tube. 2.5 μl of 1/10 ERCC spike-ins was added to each sample. Total RNA was then extracted using the hot phenol method. Briefly, 60 μl of 10% w/v SDS was added to the suspension, and the samples were mixed by inversion. Tubes were placed at 64□ for 1-2 min until clearance of the solution, then 66 μl of 3M sodium acetate solution at pH 5.2 was added and tubes were mixed by inversion. 750 μl of phenol (Roti-Aqua phenol #A980.3) was then added to each tube, mixed by inversion and incubated for 6 min at 64□. Tubes were then placed on ice to cool and spun for 15 min at 13 000 rpm, 4□. The resulting aqueous layer was transferred in a 2 ml PLG tube (5PRIME) where 750 μl of chloroform (Roth, #Y015.2) was added. After mixing by inversion, the tubes were spun for 15 min at 13 000 rpm, 4□. The obtained aqueous layer was then collected and precipitated in a 30:1 mix of 100% ethanol: 3M sodium acetate pH 6.5 at -20□ for at least 2 hr. After centrifugation for 30 min, 13 000 rpm, 4□, the pellet was washed with 70% ethanol and the air-dried pellet was resuspended in nuclease-free water. Total RNA was measured by nanodrop, and integrity was checked on TBE agarose gel. 40 μg of RNA in 39.5 μl of nuclease free water were then subjected to DNAse I treatment. Total RNA was denatured for 5 min at 65□ and put back on ice. 5 μl of DNase I (Fermentas), 5 μl of DNase I buffer (Fermentas) and 0.5 μl of Superase In (Thermo Fisher Scientific) were added to the denatured RNA and incubated at 37°C for 30 min. After incubation, 100 μl of nuclease free water was added and each reaction was placed in a PLG tube containing 150 μl of PCI. Tubes were centrifuged for 15 min at 4□, 13 000 rpm. The aqueous phases were collected and precipitated in 30:1 Ethanol/sodium acetate mix at -20□ for at least 2 hr. DNase treated pellets were collected by centrifugation (30 min, 4□, 13 000 rpm) and after 70% ethanol wash, were resuspended in 25 μl nuclease free water.

RNA-seq libraries were prepared by Vertis AG (Freising-Weihenstephan, Germany). Briefly, the ribodepleted RNA samples were fragmented using ultrasound (4 pulses of 30 s each at 4°C). Subsequently, an oligonucleotide adapter was ligated to the 3’ end of the RNA molecules. First-strand cDNA synthesis was performed using M-MLV reverse transcriptase and the 3’ adapter as primer. The first-strand cDNA was purified and the 5’ Illumina TruSeq sequencing adapter was ligated to the 3’ end of the antisense cDNA. The resulting cDNA was PCR-amplified to about 10-20 ng/μl using a high fidelity DNA polymerase. cDNA was purified with the Agencourt AMPure XP kit (Beckman Coulter Genomics) and sequenced on an Illumina HiSeq2000. Replicate 2 of the 24 minute time point for WT, Δ*proQ*, and *proQ*++ was excluded from subsequent analysis, as rRNA depletion failed.

### Processing of sequencing reads and mapping of RIF-seq data

The 75 nt RNA-seq reads were demultiplexed and quality control of each sample was performed with fastQC. Afterwards, Illumina adapters were removed with Cutadapt v4.1, and STAR (73) was used to align the reads to the SL1344 genome (NCBI accessions: FQ312003.1, HE654724.1, HE654725.1 and HE654726.1). For all analyses related to annotated genomic features such as CDSs, tRNAs, and rRNAs, gene annotations from NCBI were used. We use the same definition of transcriptional units as (54) which is based on the NCBI CDS annotations, transcription start site annotations (74), and Rho-independent terminator prediction with RNIE (75). sRNA annotations are based on (13). The ERCC92.fa sequence file for the quantification of the spike-in was obtained from ThermoScientific. For quantification, htseq-count with default options was used for counting reads aligning to CDS, sRNA and ERCC spike-ins, while the 60 base sub-genic windows were counted with the option *--nonunique all* to ensure that overlapping reads are assigned to all overlapping segments. For the 60 base windows, reads were quantified separately for the positive and the negative strand.

### Read count normalization of RIF-seq data with ERCC spike-ins

The samples were normalized using a custom implementation of the trimmed mean of M-values (76) across 30 detected ERCC spike-ins (**Fig. S2B&C**). The cutoff on the M values was set to 0.3 and the cutoff on the A values to 0.05. Only transcripts with more than 10 counts-per-million (cpm) before normalization in at least three samples in the ProQ assay were retained for further analysis.

Normalized counts-per-million (cpm) were obtained by adding a pseudo count and then dividing the read counts *Y*_*gcr*_ (*t*) by the respective library size and normalization factor *n*_*f,s*_ of the sample

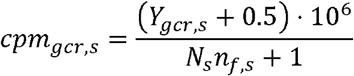

The Stan models were applied to the natural logarithm of the normalized cpm values *Y*_*gcr*_ (*t*) ≡ *In* (*cpm*_*gcr*_ (*t*)).

### Removal of batch effects: center-mean normalization

Following spike-in normalization, we observed some clustering by replicate rather than condition within time point groups (**Figure S2D&I**). To account for these batch effects, we developed a center-mean (CM) normalization procedure, which can be applied after a primary normalization, e.g. with spike-ins, and compensates for small variations in the amount of spike-ins added to the individual samples. After the normalization with spike-ins, we calculated a gene-wise mean log-count *Y*_*gc*_ (*t*) for every condition and every time-point (see **Figure S2E** for *t* = 0 min). This value was subtracted from the observed value in every sample

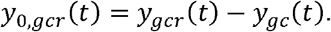

For every sample s (uniquely defined by condition *c*, time *t*, replicate *r*), we calculated the mean

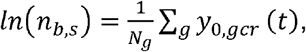

where *n*_*b,s*_ is an additional normalization constant. The batch-corrected cpm values are then given by

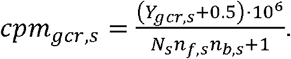

PCA confirmed that the samples separated well by time point and genotype after the CM normalization (**Figure S2F&L**), and boxplots showed an improved alignment of median logcpm values (**Figure S2G**,**H**,**J**,**K**). Before fitting the decay curves with the Bayesian models, we subtracted the mean log-count at *t* = 0 min

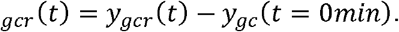

### Calculation of detection limit

In order to regularize zero counts, we have added a pseudo count of 0.5. The library sizes vary in size around 10 million reads and the normalization constants around 1. This results in an estimated minimum log-count of

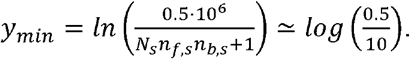

After subtracting the mean log-count at *t*= 0 min, we can calculate the detection limit for gene *g* in condition *c* as

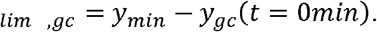

This corresponds to the minimal possible value of the log relative expression.

### Differential gene expression analysis

Log-fold changes were calculated using glmQLFit from edgeR (26) with a cutoff of 0.25 on the log-fold changes. Since batch effects were present after TMM normalization (**Fig. S1 A**,**B&E**), samples were additionally normalized using RUVg (77) (**Fig. S1C&G**). We selected the 800 least varying genes between the ΔRBP and the WT strain. Since the differences between the *proQ*++ and the WT strain were larger, we only took the 600 least varying genes between these two conditions. The intersection between these sets is 37 genes which we used as negative control (**Fig. S1F**). The number of factors of unwanted variation k was set to 6. After RUVg normalization, the samples clustered by strain (**Fig. S1C&H**). We selected differentially expressed genes at an FDR of 0.1 (**Fig. S1I-K, Table S2**).

### Extraction of RNA half-lives from RIF-seq data

We compared three statistical models, summarized in **Figure 1B**. All models assume that the normalized log counts follow a normal distribution around a condition and gene-dependent mean

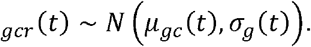

Normal distributions have been shown to effectively model RNA-seq data (78), and were computationally more efficient in our initial testing than count-based models (see the SI Appendix for a comparison between normal distributions and count-based models). The mean *μ*_*gc*_ is parameterized differently in the three statistical models:

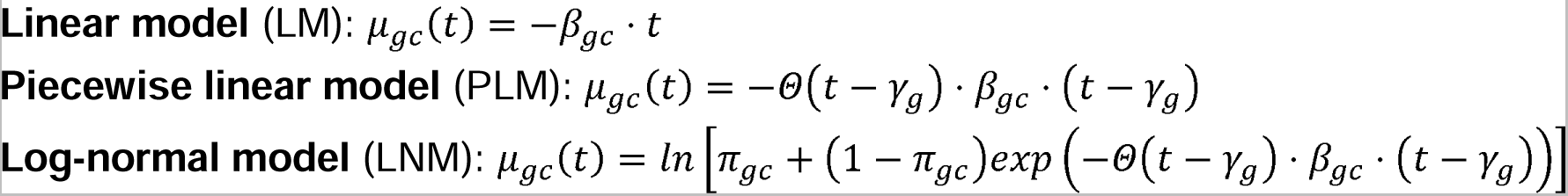

where *Θ* is the Heaviside step function which is 0 for negative arguments and 1 otherwise. The baseline parameter *π*_*gc*_ introduced in the LNM corresponds to the fraction of stable RNA for gene *g* in condition *c* as compared to steady-state levels at *t* = 0 min. Since we have no prior knowledge of the distribution of the WT decay rates, we impose a hierarchical prior with mean *μ*_*β*_ and width *σ*_*β*_ to “learn” the distribution of the decay rates and to share information across genetic loci. Additionally, we model the gene-wise standard deviation *σ*_*g*_ hierarchically, which shrinks the gene-wise variance towards the mean variance and reduces the effect of outliers. This is similar to the idea of variance shrinkage through empirical Bayes methods implemented in edgeR (26) and limma (79). Since our library sizes do not vary much between replicates within time points (Figure S2M), there were no large differences in measurement precision to account for (78). For other parameters (e.g. difference in decay rate), broad priors were chosen to minimize their influence on posterior estimates. Priors were defined as follows:

*WT* decay rate *β*_*g,WT*_ ∼ N(μ_*β*_, *σ*_*β*_)

Mutant decay rate *Δ β*_gc_ = *β*_*gc*_ - *β*_*g,WT*_ ∼ *N*(0,0.2)

Standard deviation *σ*_*g*_(*t*) ∼ *N*(*μ*_*σ*_, *σ*_*σ*_) and *σ*_*g*_ (t) ≥0

Baseline parameter *π*_*gc*_ ∼ *N* (0,0.25) and *π*_*gc*_ ∈[0,0.2]

Hyperparameters *μ*_*β*_ *μ*_σ_, *σ*_*β*_ ∼ *C* (0,1), *σ*_*σ*_∼ *N* (0.3,0.3) and *μ*_*β*_ *μ*_σ_, *σ*_*β*_ *σ*_*σ*_ ≥ 0

Elongation time *γ*_*g*_ ∼ *C*_[0,12]_(0,2)

For *π*_*gc*_ = 0, the LNM is equivalent to the PLM, which converts to the LM as *γ*_*gc*_ →0. The statistical models are fitted to the RIF-seq data using the probabilistic programming language Stan (v.2.30.1) (29) with two chains and 1000 MCMC samples each (`method=sample num_samples=1000 num_warmup=1000 adapt delta=0.95 algorithm=hmc engine=nuts max_depth=15`). The statistical model was applied to all four strains (WT, Δ*proQ, proQ*++, Δ*cspCE*) at once. The reported parameters (decay rate, half-life, transcription elongation time) correspond to the median of the 2000 MCMC samples. The median of the transcriptome-wide half-lives corresponds to the median of the 2000 MCMC samples of the hyperparameter *μ*_*β*_. In addition to the 2^nd^ replicate of the time point taken at 24 minutes for the *proQ* experiments, the 1^st^ replicate of the 4 min time point of the Δ*cspCE* mutant was removed from this part of the analysis because it clustered together with the 0 min time point (**Figure S1I**) which strongly influenced differences in decay rate in the Δ*cspCE* mutant.

### Model comparison using leave-one-out cross validation

For a quantitative comparison of the linear model (LM), the piecewise linear model (PLM) and the log-normal model (LNM), we estimated the out-of-sample predictive accuracy using leave-one-out cross validation (LOO-CV) with *Pareto-smoothed importance sampling* (PSIS) (37). The pointwise log-likelihood `log_lik` was computed in the `generated quantities` block in Stan during MCMC sampling. We used the `loo()` function from the *loo* R package (version 2.5.1), which computes the expected log-pointwise predictive density (ELPD) using PSIS.

### Calculation of transcription velocities

To calculate transcription velocities, we took advantage of ongoing transcription of RNA polymerase already bound to DNA in the RIF-seq data. We split the genome into 60 base subgenic windows, and extracted the corresponding elongation times and decay rates using the log-normal model. We split the dataset into five subsets before running the MCMC sampler (1 chain: `method=sample num_samples=1000 num_warmup=1000 adapt delta=0.95 algorithm=hmc engine=nuts max_depth=15`). Subsequently, we verified that the hierarchical parameters agreed well between the five subsets. The resulting transcription elongation times *γ* were combined with operon annotations taken from (54). We fitted a linear model with y-intercept *a*_*g*_ and slope *b*_*g*_ to the elongation times of operons or individual transcripts as shown for the mra/fts operons in **Figure S3C**, using the inverse of the 68% credible intervals of *γ*_*g*_ as obtained from the MCMC samples as weights (example: Fig. 2E). The transcription velocity *υ*_*g*_ is given by the ratio of the window size (*s*_*seg*_ = 60 nt) and the slope *b*_*g*_. Its error was calculated via error propagation 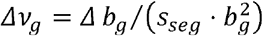. We obtain 772 operons with at least 7 nonzero segments which fulfill the quality criterium Δ *υ*_*g*_ ⁄ *υ*_g_ < 0.75.

### Calculation of Bayesian p-values

The half-lives were calculated from the decay rates *t*_1⁄2_,gc = *In* (2) ⁄ *β*_*gc*_. In order to calculate Bayesian p-values, we tested against the null hypothesis that the difference in half-life Δ*t*_1⁄2_,*c* = *t*_1⁄2_, −_*t*1⁄2,WT_ is compatible with zero. There is a limit as to how precisely we measured the WT half-lives. We determined the minimum of the 90% credible intervals of the WT half-lives (∼0.05). Assuming that we cannot measure a difference in half-life with higher precision than the WT half-lives, we selected the interval [-0.05, 0.05] as the null hypothesis. The p-value *p*_*gc*_ for gene *g* in condition *c* corresponding to the difference in decay rate *Δ t*_1⁄2,*gc*_ is given by the fraction |*S*0|⁄|*S*| of MCMC samples S= {s1,…,s_2000_} that agrees with the null hypothesis (**Figure S4A**):

For *Δ* t_1⁄2,gc_ > 0, the samples S0 = {s ∈ S|s ≦ 0.05} agree with the null hypothesis. For *Δ t*_1⁄2,g_ < 0, the samples S0 = {s ∈ S|s − 0.05}agree with the null hypothesis.

We compared the distribution of p-values to the distribution of p-values under the null hypothesis which was obtained by bootstrapping from the distribution of MCMC samples of the WT half-lives and calculating the corresponding p-values (**Figure S4B**).

### Calibration of posterior predictive p-values

In order to assign a false-discovery rate (FDR) to the p-values, we simulated a dataset with 4000 transcripts, 3 conditions (*WT, c*_1_, *c*_2_) with time points 0, 3, 6, 12, 24 and 2 conditions (*WT, c*_3_) with time points 0, 2, 4, 8, 16. We drew samples from the following distributions (which we extracted from fitting the LNM to the two RIF-seq data sets), using the definition of the LNM as given above:

Relative log-counts *ygcr* (t) ∼ *N*(*μ gc* (t),(*σ g*(t))

*WT* decay rate *β*_g,WT_ ∼ *N*(0.75,0.3)

Elongation time *γ*_g_ ∼ *C*_[0,12]_(0,2)

Standard deviation of log-counts *σ*_g_ ∼ *N*(0.35,0.23) and *σ*_g_(t) ≥ 0

Difference in decay rate *Δ β*_*gc*_ = *β*_*gc*_ − *β*_*g,WT*_ ∼ *N*(0,0.08)

Baseline parameter *π*_*gc*_ ∼ *N* (0,0.05)

Mean of relative log-counts *μ*_*gc*_ (*t*) = *In* [*π*_*gc* +_ (1-*π*_*gc*_) *exp*(− Θ(*t* − *γ*_*g*_). *β*_*gc*_. (*t* − *γ*_*g*_))].

Then, we fitted the log-normal model to this dataset. Simulated absolute differences in half-life below 0.05 (| Δ t_1⁄2,*gc*_ | ≤ 0.05) were assumed to agree with the null hypothesis. The Pearson correlation of 0.86 between simulated and fitted differences in half-life were obtained using the weightedCorr function from the *wCorr* package in R with the inverse of the size of the 90% credible intervals of the fitted half-lives as weights (**Figure S4C**). We calculated the posterior predictive p-values for the fitted differences in half-life and varied the p-value cutoff between 0 and 1 with step size 0.01. The corresponding FDR is given by the fraction of transcripts whose simulated difference in half-life agrees with the null hypothesis and the total number of transcripts with a p-value below the cutoff. Subsequently, we fitted a LOESS curve in R (`span=0.2`) to determine the FDR corresponding to any p-value cutoff (**Figure S4E, Table S1**) A p-value of 0.082 corresponds to a FDR of 0.1 which we used as a cutoff for our analysis of differentially decaying transcripts. In addition to controlling the FDR, we verified that at an FDR of 0.1, the log-normal model identifies differentially decaying transcripts with a low simulated standard deviation on log-counts *σ*_*g*_ with high sensitivity (**Figure S4D**). For this, we selected five cutoffs on standard deviation (0.05, 0.1, 0.2, 0.4, 1) and calculated the false positive rate (FPR) and sensitivity for all transcripts below the cutoff.

## Supplementary Material

**Supplementary Methods**, bacterial strains, plasmids, oligos and antibodies used in this study (**Tables S1-S4**) and Supplementary **Figures S1-14** can be found in the SI Appendix. Genome-wide *Salmonella* half-life estimates (**Dataset S1**) and log-fold changes in the absence of the global RBPs ProQ and CspC/E (**Dataset S2**), candidate lists for ProQ and CspC/E protected transcripts (**Dataset S3&4**) and significant CLIP-seq peaks for ProQ, CspC & CspE (**Dataset S5-7**) are provided as XLSX files.

## Notes

https://github.com/BarquistLab/RIF-seq_repo

### Competing Interest Statement

The authors have declared no competing interest.

### Summary of Updates

This version of the manuscript has been revised to update details of the sequencing protocol, and findings regarding the ArcA regulon have been removed.

